# An improved assembly and annotation of the allohexaploid wheat genome identifies complete families of agronomic genes and provides genomic evidence for chromosomal translocations

**DOI:** 10.1101/080796

**Authors:** Bernardo J. Clavijo, Luca Venturini, Christian Schudoma, Gonzalo Garcia Accinelli, Gemy Kaithakottil, Jonathan Wright, Philippa Borrill, George Kettleborough, Darren Heavens, Helen Chapman, James Lipscombe, Tom Barker, Fu-Hao Lu, Neil McKenzie, Dina Raats, Ricardo H. Ramirez-Gonzalez, Aurore Coince, Ned Peel, Lawrence Percival-Alwyn, Owen Duncan, Josua Trösch, Guotai Yu, Dan Bolser, Guy Namaati, Arnaud Kerhornou, Manuel Spannagl, Heidrun Gundlach, Georg Haberer, Robert P. Davey, Christine Fosker, Federica Di Palma, Andrew Phillips, A. Harvey Millar, Paul J. Kersey, Cristobal Uauy, Ksenia V. Krasileva, David Swarbreck, Michael W. Bevan, Matthew D. Clark

## Abstract

Advances in genome sequencing and assembly technologies are generating many high quality genome sequences, but assemblies of large, repeat-rich polyploid genomes, such as that of bread wheat, remain fragmented and incomplete. We have generated a new wheat whole-genome shotgun sequence assembly using a combination of optimised data types and an assembly algorithm designed to deal with large and complex genomes. The new assembly represents more than 78% of the genome with a scaffold N50 of 88.8kbp that has a high fidelity to the input data. Our new annotation combines strand-specific Illumina RNAseq and PacBio full-length cDNAs to identify 104,091 high confidence protein-coding genes and 10,156 non-coding RNA genes. We confirmed three known and identified one novel genome rearrangements. Our approach enables the rapid and scalable assembly of wheat genomes, the identification of structural variants, and the definition of complete gene models, all powerful resources for trait analysis and breeding of this key global crop. [Supplemental material is available for this article.]

---

Improvements in sequencing read lengths and throughput have enabled the rapid and cost-effective assembly of many large and complex genomes (Gnerre et al., 2011; Lam et al., 2011). Comparisons between assembled genomes have revealed many classes of sequence variation of major functional significance that were not detected by direct alignment of sequence reads to a common reference (Gan et al., 2011; 1000 Genomes Project Consortium et al., 2010; Bishara et al., 2015). Therefore, accurate comparative genomics requires that genome sequences are assembled prior to alignment, but in many eukaryotic genomes assembly is complicated by the presence of large tracts of repetitive sequences (Treangen and Salzberg, 2011; Chaisson et al., 2015) and the common occurrence of genome duplications, for example in polyploids (Blanc and Wolfe, 2004; Berthelot et al., 2014).

Recent innovations in sequence library preparation, assembly algorithms, and long-range scaffolding have dramatically improved whole genome shotgun assemblies from short read sequences. These include PCR-free library preparation to reduce bias (Aird et al., 2011), longer sequence reads, and algorithms that preserve allelic diversity during assembly (Weisenfeld et al., 2014). Shortread assemblies have been linked into larger chromosome-scale scaffolds by Hi-C in vivo (Lieberman-Aiden et al., 2009) and in vitro (Putnam et al., 2016) chromatin proximity ligation, and by linked-read sequencing technologies (Mostovoy et al., 2016; Weisenfeld et al., 2016). Although it is more expensive than short read sequencing approaches, Single Molecule Real Time (SMRT) sequencing improved the contiguity and repeat representation of mammalian (Pendleton et al., 2015; Gordon et al., 2016) and diploid grass genomes (Zimin et al., 2016). SMRT technologies are also being used to generate the complete sequence of transcripts, increasing the accuracy of splicing isoform definition (Abdel-Ghany et al., 2016).

The assembly of the 17Gbp allohexaploid genome of bread wheat (*Triticum aestivum*) has posed major difficulties, as it is composed of three large, repetitive and closely related genomes (Moore et al., 1995). Despite progressive improvements, an accurate and near-complete wheat genome sequence assembly and corresponding high-quality gene annotation has not yet been generated. Initial whole genome sequencing used orthologous Poaceae protein sequences to generate highly fragmented gene assemblies (Brenchley et al., 2012). A BAC-based assembly of chromosome 3B provided major insights into wheat chromosome organization (Choulet et al., 2014). Illumina sequencing and assembly of flowsorted chromosome arm DNA (Chromosome Survey Sequencing, CSS) identified homoeologous relationships between genes in the three genomes, but the assemblies remained highly fragmented (IWGSC, 2014). Recently a whole genome shotgun sequence of hexaploid wheat was assembled and anchored, though not annotated, using an ultra-dense genetic map (Chapman et al., 2015). The assembly contained ~ 48.2% of the genome with contig and scaffold N50 lengths of 8.3kbp and 25kbp, respectively.

Here we report a new sequence assembly and annotation of the allohexaploid wheat landrace Chinese Spring (CS42). Our approach is open source, rapid and scalable, and has enabled a more in-depth analysis of sequence and structural variation in this key global crop. The scaffolds cover 13.4Gbp of the genome with an N50 of 88.8kbp, and are classified into chromosome arms. The annotation, supported by extensive transcriptome sequence and 1.5 million full length SMRT cDNA sequences, contains 104,091 high confidence protein-coding genes.

## Results

### DNA library preparation and sequencing

We reduced bias and retained maximum sequence complexity by using unamplified libraries for contig generation (Kozarewa et al., 2009) and precisely sized mate-pair libraries for scaffolding (Heavens et al., 2015). Libraries were sequenced using Illumina pairedend (PE) 250bp reads to distinguish closely related sequences. In total, 1.1 billion PE reads were generated to provide 33 × sequence coverage of the CS42 genome (Supplemental Table S4.1). For scaffolding, long mate-pair (LMP) libraries with insert sizes ranging from 2480 to 11,600bp provided 53 × sequence coverage, and Tight, Amplification-free, Large insert PE Libraries (TALL) with an insert size of 690bp provided 15 × sequence coverage (Supplemental Table S4.2).

### Genome assembly

Nearly 3 million contigs (of length greater than 500bp) were generated using the w2rap-contigger (Clavijo, 2016) with an N50 of 16.7kbp (Supplemental Table S4.3). After scaffolding using SOAPdenovo (Luo et al., 2012), the assembly contained 1.3 million sequences with an N50 of 83.9kbp. To generate the TGACv1 assembly, scaffolds were classified to chromosome arms using raw CSS reads (IWGSC, 2014) and subsequently screened with a two-tiered filter based first on their length and their k-mer content (see Supplemental Information S4.5). The approach removed short, redundant sequences from the assembly minimising the loss of unique sequence content, leading to an increase in scaffold N50 to 88.8kbp.

The genome of a synthetic wheat line W7984 was previously assembled with an improved version of meraculous (Chapman et al., 2011) using 150bp PE libraries with varying insert sizes, for a combined genome coverage of 34.3 ×, together with 1.5kbp and 4kbp LMP libraries for scaffolding (Chapman et al., 2015). This contig assembly, with an N50 of 8.3kbp, covered 8Gbp of the genome while the scaffold assembly covered 8.21Gbp with an N50 of 24.8kbp. In comparison, the TGACv1 assembly represents almost 80% of the 17Gbp genome, a 60% improvement in genome coverage. The contiguity of the TGACv1 assembly is nearly four times that of the W7984 assembly and thirty times that of the CSS assembly (Table 1; IWGSC (2014)).

**Table 1:** Comparison of TGACv1 scaffolds to the IWGSC and Chapman assemblies of hexaploid wheat. Numbers are calculated using sequences greater than 500bp and including gaps (Ns) for each assembly.

|  | Size (Gb) | Seq. count | N20 (kb) | N50 (kb) | N80 (kb) | %Ns | % of genome |
| --- | --- | --- | --- | --- | --- | --- | --- |
| TGACv1 | 13.43 | 735,943 | 180.1 | 88.8 | 32.8 | 5.7 | 78.8 |
| W7984 | 8.21 | 955,122 | 47.1 | 24.8 | 9.9 | 15.2 | 48.2 |
| CSS | 8.32 | 4,061,833 | 8.6 | 3.3 | 1.2 | 1.0 | 48.9 |

The KAT spectra-cn plot generated from TGACv1 (Figure 1A) showed that k-mers found at low frequency (<12), representing sequencing errors, were not found in the assembly (shown by the black distribution at k-mer multiplicity <12). Most sequence content was represented in the assembly once (shown by the main red distribution), with k-mers originating from the repetitive and the homoeologous regions of the genome represented at higher frequencies (>50). This indicated a low level of chimeric assemblies and established the accuracy of the assembly and scaffolding methods. K-mer spectra analysis of the CSS assembly (Figure 1B), revealed a larger fraction of absent content, corresponding to the black distribution between k-mer multiplicity 15 and 45, and greater amounts of duplication in the single copy regions of the assembly, corresponding to the purple and green areas of the main red distribution. A large amount of content in the CSS assembly did not appear as sequenced content of the reads in our PCR-free paired end data, as shown in the red bar at k-mer multiplicity equal to 0, indicating a high level of chimeric sequences or consensus problems.

**Figure 1:**
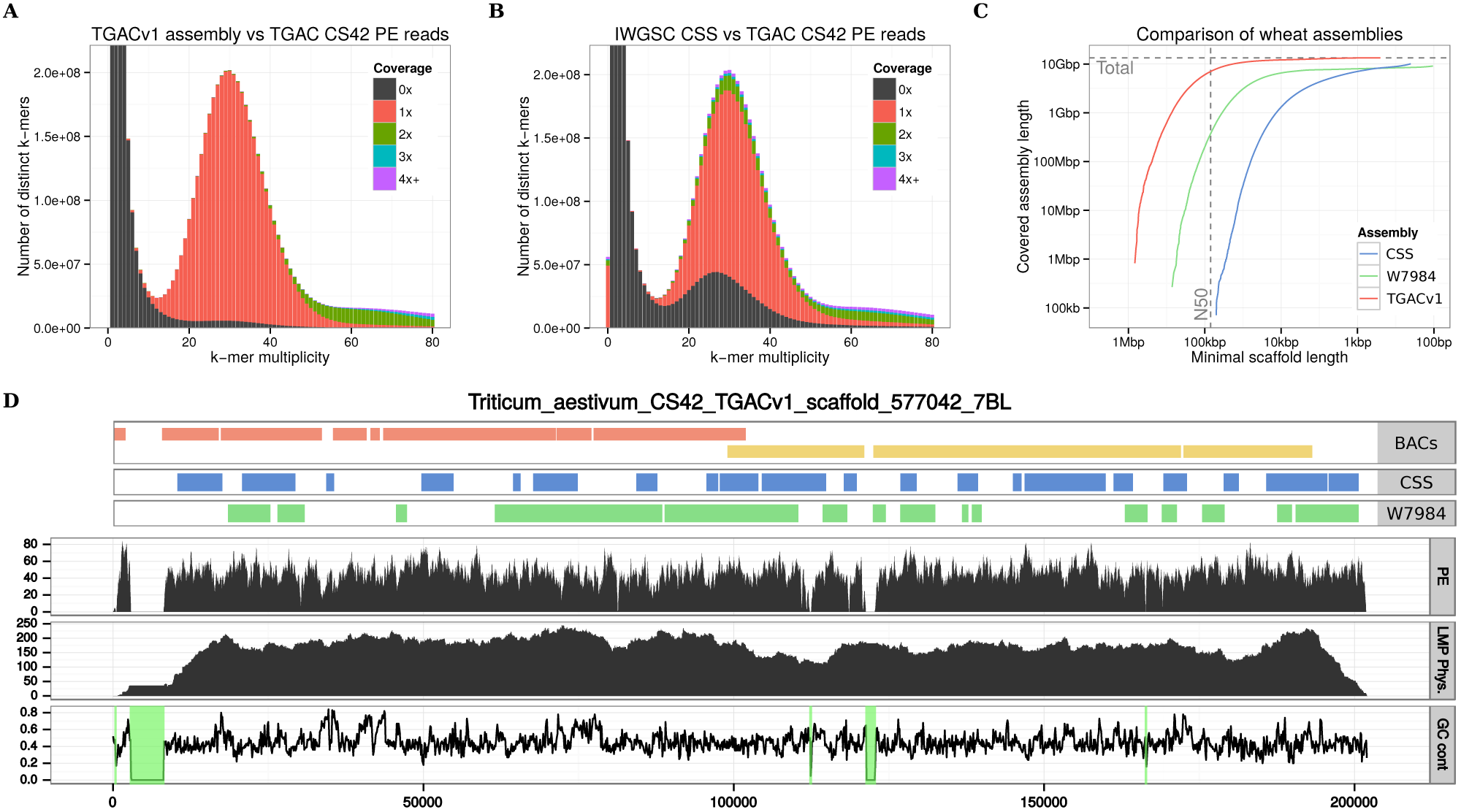
Summary of the TGACv1 wheat genome sequence assembly. **A,B**: KAT spectra-cn plots comparing the PE reads to the TGACv1 scaffolds (**A**) and CSS scaffolds (**B**). Plots are coloured to show how many times fixed length words (k-mers) from the reads appear in the assembly; frequency of occurrence (multiplicity, x-axis) and number of distinct k-mers (y-axis). Black represents k-mers missing from the assembly, red represents k-mers that appear once in the assembly, green twice, etc. Plots were generated using k = 31. The black distribution between k-mer multiplicity 15 and 45 in (B) represents k-mers that do not appear in the CSS assembly. **C:** Comparison of scaffold lengths and total assembly sizes of the TGACv1, W7984, and CSS assemblies. **D:** Scaffold 577042 of the TGACv1 assembly. Tracks from top to bottom: aligned BAC contigs, CSS contigs, W7984 contigs, coverage of PE reads, coverage of LMP fragments, and GC content with scaffolded gaps (N stretches) with 0% GC highlighted in green. There are two BACs (comprised of 7 and 4 contigs each), 22 CSS contigs, and 15 W7984 contigs across the single TGACv1 scaffold.

90.7% of the genes previously identified on the 3B BAC-based assembly (Choulet et al., 2014) aligned to TGACv1 scaffolds (Table 2), compared with 68–74% aligned to W7984 and 67.9% aligned to CSS chromosome 3B scaffolds. This demonstrated both the improved representation of the TGACv1 assembly and the precision of chromosome classifications.

**Table 2:** Comparison of TGACv1 chromosome 3B scaffolds to BACbased scaffolds (Choulet et al., 2014), and 3B scaffolds from the W7984 and CSS assemblies. Numbers are calculated using sequences greater than 500bp and including gaps (Ns) for each assembly.

|  | Scaffold count | N50 (kb) | Total seq. (Mb) | Gene count | % genes |
| --- | --- | --- | --- | --- | --- |
| 3B ref. | 2,808 | 892.4 | 832.8 | 7,703 | 100.0 |
| TGACv1 | 29,090 | 116.5 | 790.0 | 6,983 | 90.7 |
| W7984 | 26,206 | 30.6 | 479.4 | 5,671 | 73.6 |
| CSS | 272,072 | 3.4 | 557.2 | 5,233 | 67.9 |

Alignment of TGACv1 3B scaffolds to the 3B BAC-based pseudomolecule (Figure 2A,C) showed that they were largely in agreement. Two examples of apparent disagreement are shown in Figure 2B,D. Scaffold_221671_3B spanned a gap of 700kbp in the 3B BAC assembly, and re-oriented and removed a duplication, by identifying both ends of a CACTA element (Figure 2B). Scaffold_ 220592_3B spanned 582kbp and diverged in one location (Figure 2D), and contained a Sabrina solo-LTR with a characteristic ATCAG target site duplication (TSD). In scaffold_220592_3B the TSD was present on either side of the Sabrina_3231 element, while in the BAC-based scaffold Sabrina homology ended in Ns. In the BAC-based assembly only one side of the disjunction showed alignment similarity to CACTA_3026, which was found to be complete in scaffold_220592_3B and spanned the disjunction (Figure 2D). These two examples illustrate how the TGACv1 assembly generated accurate scaffolds spanning typical complex and long tracts of repetitive DNA characterising the wheat genome, which were misassembled in the BAC-based approach.

**Figure 2:**
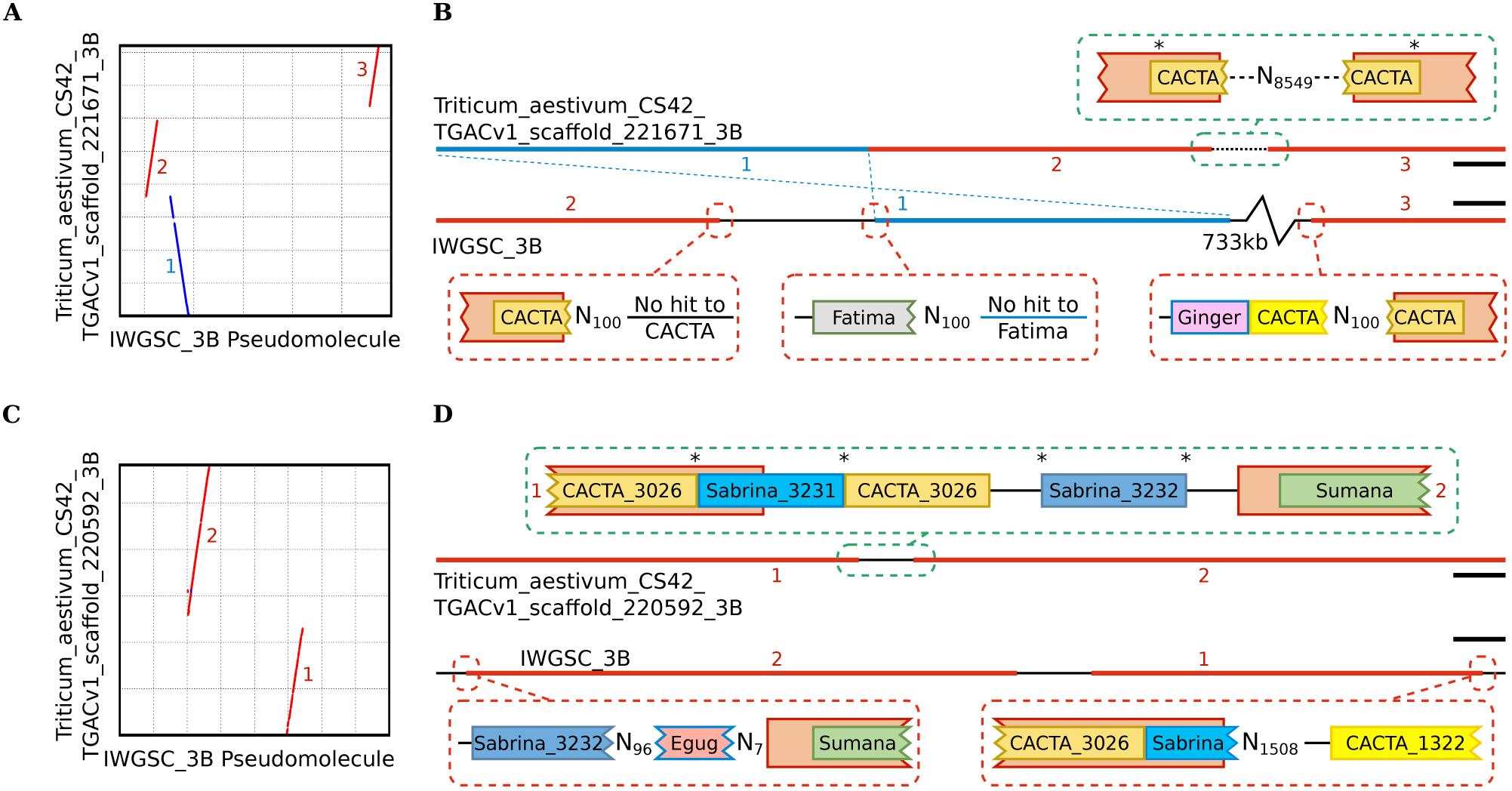
Comparative alignment of TGACv1 scaffolds with the 3B BAC-based pseudomolecule. **A,C**: Dot plots between TGACv1 scaffolds and 3B show disruptions in sequence alignment including rearrangements (red) and inversions (blue). **B,D**: Graphical representation of sequence annotations in disrupted regions. Junctions in the TGACv1 scaffolds are consistent with a complete retroelement spanning the junction that includes identical TSD on either side of the retroelement (asterisks). Corresponding regions in the 3B BAC-based pseudomolecule are characterised by Ns that produce inconsistent alignment of retroelements across putative junctions. Retroelements of the same family (CACTA, Sabrina) but matching distinct members in the TREP database are indicated by different colours. Numbers adjacent to sequences correspond to regions shown in panel A and C, respectively. Scale bars correspond to 10kbp (**B**) and 30kbp (**D**).

### Repetitive DNA composition

Supplemental Table S7.1 shows the class I and class II mobile element composition of wheat based on the TGACv1 assembly. More than 80% of the 13.4Gbp assembly was composed of approximately 9.7 million annotated transposable element entities, of which nearly 70% were retroelements (class I) and 13% DNA transposons (class II). Among the class I elements, Gypsy and Copia LTR retroelements comprised the major component of the repeats, while CACTA DNA elements were highly predominant among class II DNA repeat types. No major differences in the repeat composition of the three genomes were apparent. Compared with Brachypodium distachyon, which has a related but much smaller genome (Vogel et al., 2010), there has been a greater than 100 × increase in repeat content, driven by both class I and class II expansion. The preponderance of CACTA DNA elements in the wheat genome emerged during this massive expansion.

### Gene prediction and annotation

A total of 217,907 loci and 273,739 transcripts were identified from a combination of cross-species protein alignments, 1.5 million high-quality long PacBio cDNA reads, and over 3.2 billion RNA-Seq read pairs covering a range of tissues and developmental stages (Table 3, Supplemental Information S8).

**Table 3:**
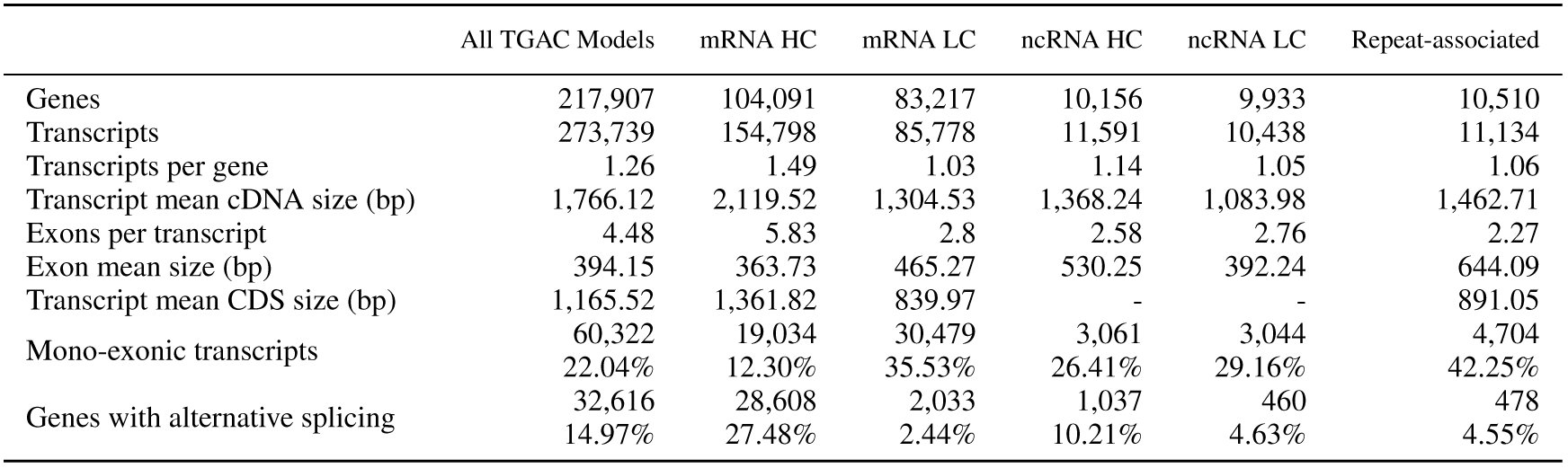
Characteristics of predicted high (HC) and low (LC) confidence wheat genes including coding (mRNA) and long non-coding (ncRNA) RNA.

Loci were identified as coding, long ncRNA, or repeat associated, and were classified as high or low confidence based on similarity to known plant protein sequences and supporting evidence from wheat transcripts (Supplemental Information S8.5.5). We assigned 104,091 coding genes (154,798 transcripts) as high confidence (HC), of which 95,827 spanned at least 80% of the length of the best identified homolog (termed protein rank 1, P1, in the annotation; Supplemental Figure S8.1 and Supplemental Information S8.5.1). The HC protein-coding set contained 51,851 genes confirmed by a PacBio transcript (Transcript rank 1, T1) and an additional 29,996 genes fully supported by assembled RNASeq data (T2), providing full transcriptome support for 81,847 (78.63%) HC genes. Gene predictions were assessed by identifying 2707 single copy genes common to *B. distachyon, Oryza sativa, Sorghum bicolor, Setaria italica* and *Zea mays*. A single orthologous wheat gene was identified for 2686 (99.22%) of these, with 2665 (98.45%) classified as HC and 21 (0.78%) in the low confidence (LC) set. A high coherence in gene length (r=0.969) was found between wheat and *B. distachyon proteins* (Supplemental Figure S8.2). These findings show that the HC gene set is robust and establishes a lower bound estimate for the total number of protein-coding genes in wheat. An additional 103,660 loci were defined as LC (i.e. gene models with all their transcripts either having less than 60% protein coverage or lacking wheat transcript support). These include bona fide genes that were fragmented due to breaks in the current assembly, wheat specific genes, and genes without transcriptome support Supplementary Table S8.8).

We also identified 10,156 HC non-coding genes with little similarity in protein databases and low protein coding potential. The majority of these genes are located in intergenic regions (8854, or 87.18%), while most of the remaining 1302 are antisense to coding genes (1082, or 10.65%; see Supplementary Information S8.5.8). 5413 of wheat non-coding genes (53.30%) were detected in at least one, of the two sequenced wheat diploid progenitor species *T. urartu* and *Ae. tauschii* (at least 90% coverage and 90% identity; see Supplementary Information S8.5.8).

To obtain additional support for gene predictions, a proteome map was constructed from 27 wheat tissues (Supplemental Information S9). This identified 2,106,323 significant peptide spectrum matches corresponding to 102,379 distinct peptides. Of these, 96.20% matched HC genes, while 13.29% were assigned to LC genes. For 56,391 genes (43,431 HC, 12,960 LC) we were able to identify at least one peptide confirming the predicted coding sequence. Due to the hexaploid nature of wheat, only 22.1% of the peptides could be assigned to a single gene. Applying progressively stricter filters, by requiring at least 2 or 5 peptides, confirmed the protein sequence of 30,607 and 17,316 HC genes, respectively. 10,819 genes met the criteria of having support from multiple peptides with at least one uniquely identifying peptide, and were considered as unambiguously corroborated by proteomic data. Among the LC genes, only 368 were identified by two or more peptides that did not match any HC gene, further supporting confidence assignments. Among these, 343 were classified as LC due to having less than 60% the length of the identified homolog, while the remaining 25 genes were classified as LC due to either repeat association or lack of wheat transcript support.

We compared the TGACv1, CSS (IWGSC, 2014) and chromosome 3B (Choulet et al., 2014) gene models. Of the 100,344 HC genes in the IWGSC annotation (PGSB/MIPS version 2.2 and INRA version 1.0 from Ensembl release 29) we were able to transfer 97,072 (97%) to the TGACv1 assembly with stringent alignment parameters (at least 90% coverage and 95% identity). Fewer (72%) of IWGSC (IWGSC, 2014) low confidence, unsupported, repeat associated and non-coding loci could be aligned (at least 90% coverage and 95% identity) likely reflecting differences between the assemblies of repeat rich and difficult to assemble regions. Of the TGACv1 HC genes, 61% overlapped with an aligned IWGSC HC gene and 78% to the full IWGSC gene set (Supplemental Information S8.5.7). Less agreement was found between TGACv1 LC and ncRNA genes and the IWGSC annotation, with only 8% overlapping IWGSC HC loci and 40% overlapping the full IWGSC gene set (Figure 3A). Of the 22,904 (22%) high confidence TGACv1 genes not overlapping a transferred IWGSC gene, 19,810 (86%) had cross species protein similarity support with 6665 (29%) fully supported by a PacBio transcript (Figure 3A). We identified 13,609 TGACv1 genes that were overlapped by transcripts originating from two or more IWGSC genes in our annotation, indicating that they were likely fragmented in the CSS assembly. In 8175 of these cases (60%) we were able to find a PacBio read fully supporting our gene model. These differences reflect improvements in contiguity, a more comprehensive representation of the wheat gene space in our assembly, and improved transcriptome support for annotation.

**Figure 3:**
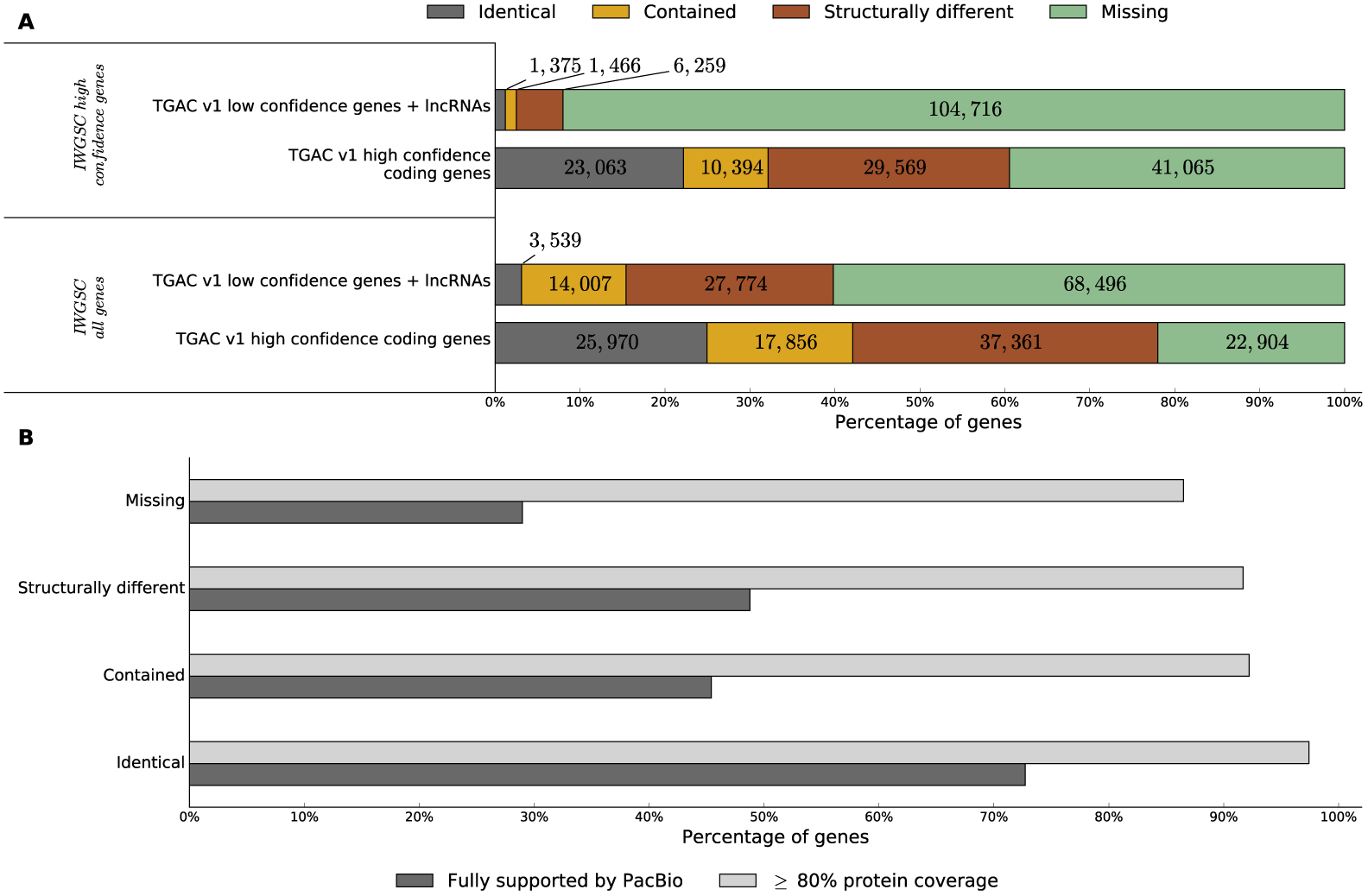
Comparison between IWGSC annotation and TGACv1 high and low confidence genes. IWGSC genes were aligned to the TGACv1 assembly (gmap, ≥ 90% coverage, ≥ 95% identity) and classified based on overlap with TGACv1 genes. **A**: Identical - shared exon-intron structure; Contained - exactly contained within the TGACv1 gene; Structurally different - alternative exon-intron structure; Missing - no overlap with IWGSC. **B**: Bar plot showing proportion of high confidence TGACv1 protein coding genes supported by protein similarity or PacBio data, genes are classified based on overlap with the full set of IWGSC genes.

### Alternative splicing

Alternative splicing is an important mRNA processing step that increases transcriptome plasticity and proteome diversity (Staiger and Brown, 2013). The TGACv1 annotation includes high-quality alternative splicing variants identified from PacBio transcriptome reads. To provide a more comprehensive representation of alternative splicing we subsequently integrated transcript assemblies generated from six strand-specific Illumina libraries (Supplementary Information S8.6 and Table S8.1). This added a further 121,997 transcripts, increasing the number of genes with splice variants from 15% in the TGACv1 annotation to 31% in the supplemented set of transcripts (i.e. incorporating Illumina RNA-Seq assemblies), and increasing the average number of transcripts per gene from 1.26 to 1.88. When considering only HC genes, the number of alternatively spliced genes was increased from 27.48% to 48.80% (2.36 transcripts per gene), similar to that observed in a wide range of plant species (Zhang et al., 2015).

Intron retention (IR) was the prevalent alternative splicing event in wheat (34%) followed by alternative 3’ splice sites (A3SS; 27%), exon skipping (ES; 20%), alternative 5’ splice sites (A5SS; 19%) and mutually exclusive exons (MXE; 0.04%). This was similar to previous analyses of chromosome 3B (Pingault et al., 2015), and IR is also predominant in barley (Panahi et al., 2015). Alternative splicing coupled to nonsense mediated decay (NMD) regulates gene expression (Lykke-Andersen and Jensen, 2015). We found 22% of all transcripts (17% of all genes) and 29% of multi-exonic HC protein coding transcripts (33% genes) may be potential targets for NMD. Intron retention was the most common splicing event leading to NMD sensitivity, with 40% of IR transcripts identified as potential NMD targets (34% ES, 38% A5SS, 34% A3SS, 26% MXE). This suggests a potentially substantial role for alternative splicing / NMD in regulating gene expression in wheat.

### Gene families

HC and LC gene families were analysed separately using OrthoMCL version 2.0 (Li (2003); Supplemental Figures S10.1 and S10.2). Splice variants were removed from the HC gene data set, keeping the representative transcript for each gene model (see Supplemental Information S8.5.6 and S10.1), and datasets were filtered for premature termination codons and incompatible reading frames. For the HC gene set, a total of 87,519 coding sequences were clustered into 25,132 gene families. The vast majority of HC gene families contained members from the A, B and D genomes, consistent with the relatively recent common ancestry of the A and B genomes and the proposed hybrid origin of the D genome from ancestral A and B genomes (Marcussen et al., 2014). Subsets of gene families and singleton genes (those not clustered into any family) were classified to identify a) genes and families that are A, B or D genome-specific, b) gene families with expanded numbers in one genome, and c) wheat gene families that are expanded relative to other species. These gene sets were analysed for over-represented Gene Ontology (GO) terms, shown in Supplementary File **S2**. Gene families that were significantly expanded in wheat compared to *Arabidopsis, rice, sorghum* and *Brachypodium* include those encoding proteins involved in chromosome maintenance and reproductive processes, and protein and macromolecule modification and protein metabolism processes. The D genome has expanded gene families encoding phosphorylation, phosphate metabolism and macromolecule modification activities, while the B genome has expanded gene families encoding components of chromosome organisation, DNA integration and conformation/unwinding, and telomere maintenance. The B genome is derived from the *Sitopsis* section of the Triticeae, which has contributed genomes to many polyploid Triticeae species (Riley et al., 1961), suggesting B genomes may have contributed gene functions for establishing and maintaining polyploidy in the Triticeae. This is supported by the location of the major chromosome pairing *Ph1* locus on chromosome 5B (Griffiths et al., 2006).

### Genome organization

A corrected version of the POPSEQ genetic map (Chapman et al., 2015) was used to order TGACv1 scaffolds along chromosomes (Supplemental information S5). This uniquely assigned 128,906 (17.5%) of the 735,943 TGACv1 scaffolds to 1051 of 1187 genetic bins (class 1, Supplemental Information S5) to form the final TGACv1 map. The total length of these scaffolds is 8,551,191,083bp, representing 63.68% of the TGACv1 assembly and 50.52% of the 17Gbp wheat genome. The TGACv1 map includes 3927 (3.05%) scaffolds that were not previously assigned to a chromosome arm (Supplemental Information S5) and 380 (0.295%) scaffolds whose CSS-based chromosome assignment disagrees with its position according to the TGACv1 map. A further 13,019 (1.77%) scaffolds were ambiguously assigned to different cM positions on the same chromosome (class 2), 489 (0.07%) scaffolds were assigned to homoeologous chromosomes (class 3), and 3320 (0.45%) scaffolds had matching markers with conflicting bin assignment (class 4). The TGACv1 map encompasses 38,958 of the 53,792 scaffolds containing at least one annotated HC protein-coding gene (72.42%), comprising gene sequences of 307,085 968bp (73.28% of total predicted gene sequence space). In total, we were able to assign genetic bins to 75,623 (72.65%) of the HC genes.

Chromosomal locations of related genes were identified by anchoring to the TGACv1 map and are displayed in Figure 4. Analysis of OrthoMCL outlier triads (Supplemental Information S6 and S10) provided genomic support for known ancestral reciprocal translocations between chromosome arms 4AL and 5AL, a combination of pericentromeric inversions between chromosome arms 4AL and 5AL, and a reciprocal exchange between chromosome arms 4AL and 7BS (Devos et al., 1995). Several putative novel chromosomal translocations were also identified (Figure 4 and Supplemental File S3). As these may have originated in the parental lines used in the POPSEQ map rather than in CS42, nine genes in the predicted translocations (6 previously known and 3 novel) were tested using PCR assays on Chinese Spring chromosomal deletion stocks (Sears, 1966). Three known translocation events, 4AL-5AL and 4AL-7BS (Devos et al., 1995) and 5AL-7BS (Ma et al., 2013), and one previously unidentified translocation 5BS-4BL were validated by PCR assays.

**Figure 4:**
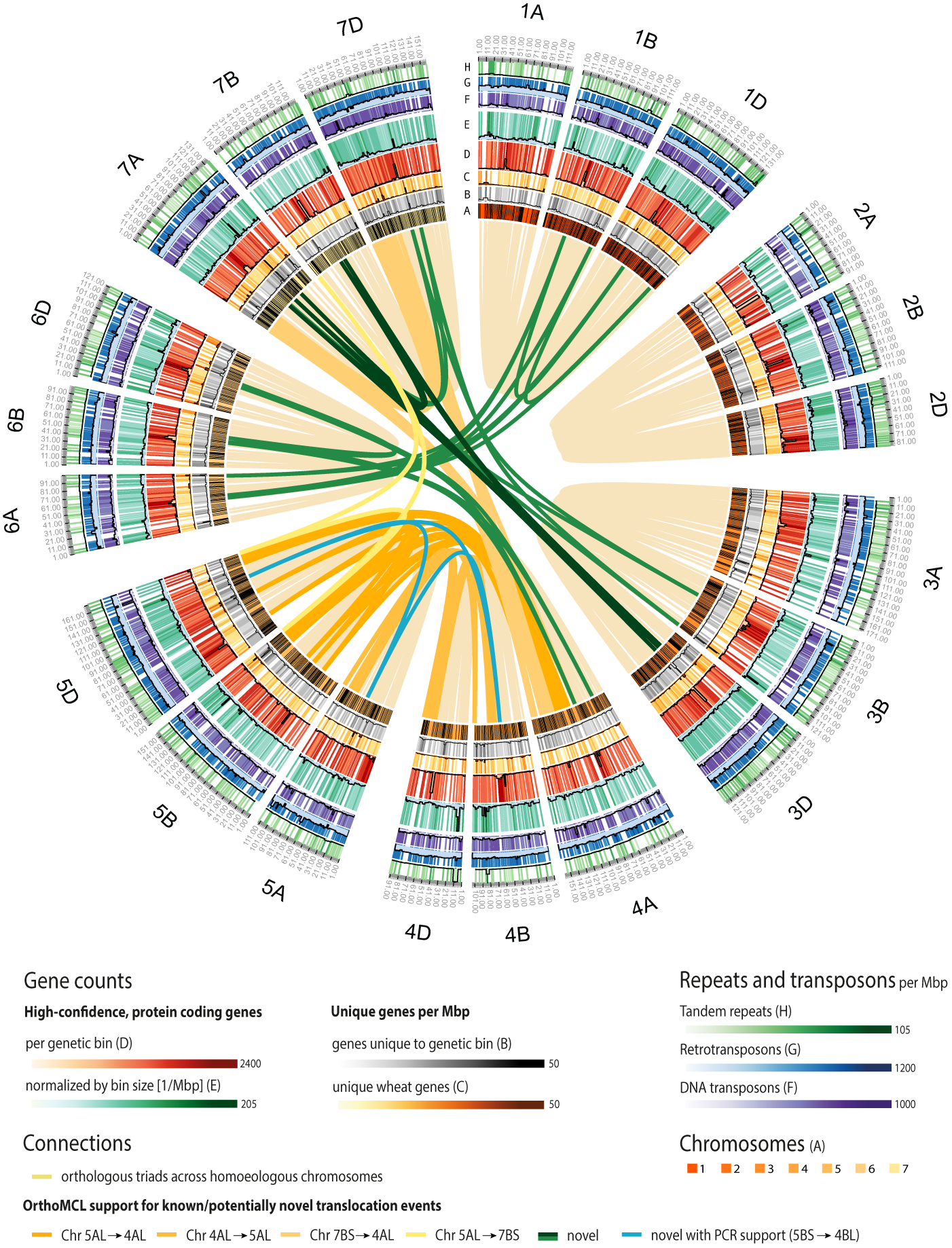
Circular representation of the TGACv1 CS42 assembly. Chromosomes, genetic bins, and genomic features are visualised on the outer rings (**A-H**) and interchromosomal links identify known and potentially novel translocation events. (**A**) The seven chromosome groups of the A, B and D genomes, scaled by number of genetic bins (black bands). (**B-H**) Combined heatmap/histogram representations of genomic features per genetic bin. With the exception of (**D**) all counts are normalised by the size of the genetic bin in Mbp, calculated as the total size of scaffolds assigned to the bin. (**B**) Distribution of unique genes, i.e. genes that did not have orthologs in a genome-wide OrthoMCL screen. (**C**) Distribution of wheat-specific genes (**D-E**) Number of HC protein-coding genes. (**F**) Distribution of DTC, DTM, and DTH DNA transposons (Supplemental Table S7.1). (**G**) Distribution of RLX, RLC, RLG, RXX, and RIX retrotransposons. (**H**) Distribution of tandem duplications. Light yellow links connect homoeologous OrthoMCL triads. Dark yellow-coloured links connect genetic bins harboring OrthoMCL outlier triads (Supplemental Information S6) that identify known translocation events. Dark green links connect genetic bins harbouring at least 3 OrthoMCL outlier triads that may support novel translocation events. The cyan link shows a novel PCR-validated translocation event between chromosomes 5BS-4BL.

### Gene expression

To explore global gene expression patterns we mapped multiple wheat RNA-Seq datasets to the TGACv1 transcriptome (Supplemental Table S11.1). Seventy-five percent of RNA-Seq reads mapped to the TGACv1 transcriptome (Supplemental Table S11.1), and 78% of the HC protein coding transcripts were expressed above the background level of 2 tpm (Wagner et al., 2013). Interestingly, 23% of the LC genes were also expressed above 2 tpm. Expression levels of genes across chromosomes were similar, with the exception of 19 genetic bins that had increased expression (defined as “hotspots” with a median expression level greater than 20 tpm, containing on average 5 genes) across the six tissues examined (Supplemental Figure S11.1). Hotspots tended to be enriched for genes encoding components of the cytoskeleton, ribosome biogenesis, and nucleosome assembly that were expressed at high levels in all tissues. Other notable hotspots were enriched in genes of photosystem I formation in leaf tissues, and nutrient reservoir activity in seed tissues.

The more complete and accurate annotation provided an opportunity to analyse patterns of transcript levels in homoeologous triads. Transcript levels of 9642 triads were analysed in response to biotic and abiotic stress using publicly available RNA-Seq (Supplemental Table S11.2), selected as they all used 7-day old seedlings, were replicated, and assessed dynamic transcriptional responses to standardised treatments. Across treatments, 26% (2424 of 9159) of expressed triads showed higher expression in one or two genomes in at least one stress condition (rather than balanced expression of three genomes; see Supplemental Information S11.5). Abiotic stress led to more differentially-regulated transcripts, compared to biotic stress responses, across all three genomes. To assess the conservation of this stress response between homoeologs, we classified each homoeolog as either up-regulated (greater than 2-fold change, UP), down-regulated (less than 0.5-fold change, DOWN) or flat (between 0.5 – 2 - fold change). We then assessed whether the individual homoeolog response to stress compared to control conditions was consistent (Supplemental Table S11.3). 80% (± 5.1% SE) of triads were not differentially expressed in response to the stress treatments and were excluded from further analysis. The most frequent pattern of differential triad expression was a single homoeolog UP or DOWN, with the other two remaining flat (Figure 5; 79–99% across conditions). Triads in which either all homoeologs were expressed in the same pattern (“3 UP” or “3 DOWN”) were rare, as were triads in which homoeologs were expressed in opposite directions. This is consistent with Liu et al. (2015), who identified between 13% and 41% of homoeolog triads in which homoeologs did not respond to the same degree in response to stress conditions.

**Figure 5:**
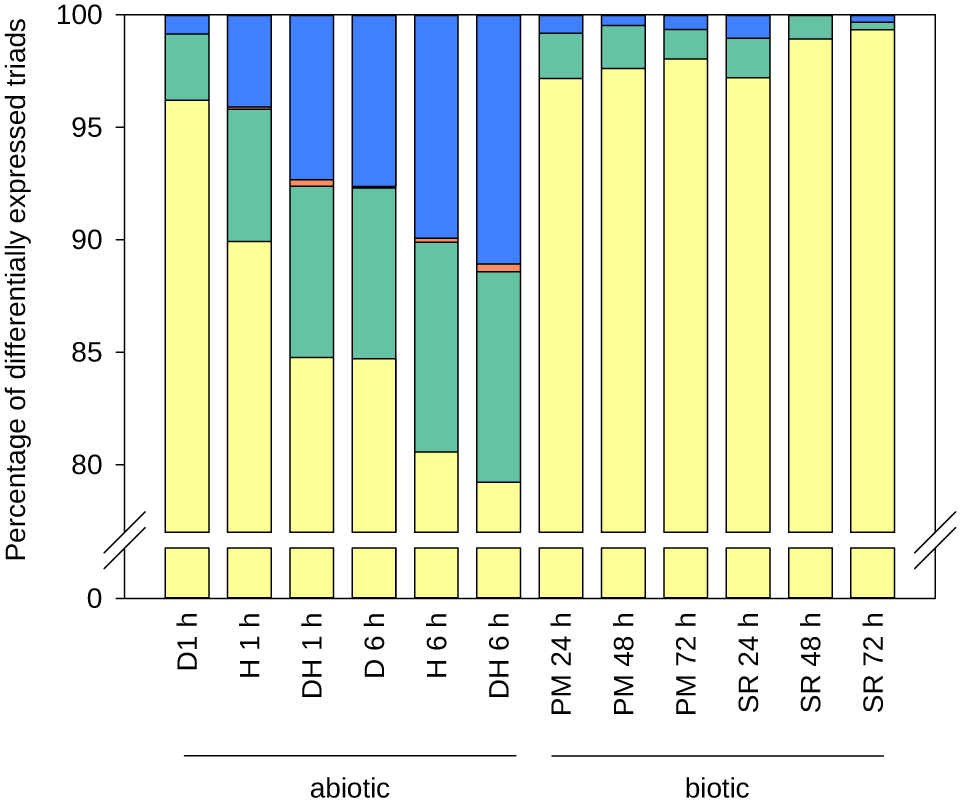
Response of differentially expressed (DE) triads to stress treatments according to the number and pattern of DE homoeologs. Triads were classified as having one homoeolog DE (yellow), two homoeologs DE with same direction of change (green), three homoeologs DE with same direction of change (orange), or opposite direction of change between DE homoeologs (blue). The stresses applied were drought (D), heat (H), drought and heat combined (DH), powdery mildew (PM) and stripe rust (SR), with the duration of stress application indicated in hours (h).

The genomic context of differences in homoeolog expression was explored in genomic regions containing at least five high confidence genes in syntenic order on all three genomes, of which at least one homoeolog was expressed over background levels in root, shoot and endosperm tissue at 10 and 20 days post anthesis (DPA) (Supplemental Table S11.1, DRP000768 and ERP004505; Pfeifer et al. (2014)). Of the four blocks meeting these criteria, one showed equal expression of all fifteen homoeologs in at least one of the tissues, while the other three blocks showed unbalanced expression of at least one homoeolog (Supplementary Figure S11.3). All blocks exhibited major structural and promoter sequence differences, and variant transcription start sites (Supplementary Figure S11.3). These multiple types of genomic differences all have the potential to contribute to unbalanced expression. To facilitate further expression studies the expression atlas at http://www.wheat-expression.com has been updated with the TGACv1 annotation and expression data from 424 RNA samples (Borrill et al., 2016).

### Gene families of agronomic interest

#### Wheat disease resistance genes

Plant disease resistance (R-) genes termed Nucleotide Binding Site-Leucine Rich Receptors (NBS-LRRs; Dodds and Rathjen (2010)) are challenging to assemble as they are often organized in multi-genic clusters with many tandem duplications and rapid pseudogenisation. The TGACv1 assembly contains 2595 NBS-containing genes (Table 4) of which 1185 are NBS-LRR genes. Among these, 98% have complete transcripts compared to only 2% in the CSS assembly. We also used NLR-parser (Steuernagel et al., 2015) to predict the coiled-coil (CC-) NBS-LRR subclass of R - genes. We identified 859 complete CC-NBS-LRR genes supported by specific MEME motifs (Jupe et al., 2012), compared to 225 in the CSS assembly (Table 4). The total of 1185 wheat NBS-NLRs was consistent with that found in diploid wheat progenitors (402 NLRs in *T. urartu*) and diploid relatives (438 in O. *sativa*; Sarris et al. (2016)). Nearly 90% of CS42 R-genes were unambiguously assigned to chromosome arms and 57% (674/1185) were anchored to the TGACv1 map. The number of R-genes per scaffold ranged from 1 to 31, compared to only 2 to 3 R-gene per scaffold in the CSS wheat assembly (IWGSC, 2014). This finding is corroborated by BAC sequence assemblies (Supplementary Figure S12.1).

**Table 4:** Disease Resistance and Gluten gene repertoires in the TGACv1 assembly. Resistance genes were identified by their characteristic domain architecture (Sarris et al., 2016). Gluten genes were identified by sequence similarity to either a gliadin, glutenin, or generic prolamin class, representing prolamin-like glutens discovered in oat (avenin), wheat (farinin), or barley (hordein). See Supplementary Information, Section 12.

| R-genes |  |  | Gluten genes |  |  |
| --- | --- | --- | --- | --- | --- |
|  | CSS | TGACv1 |  | CDS | Pseudogenes |
| *NBS-containing (Pfam)* | 1,224 | 2,595 | *Gliadins* |  |  |
| fragmented | 1,188 | 65 | alpha | 29 | 9 |
| complete transcript | 36 | 2,530 | gamma | 18 | 0 |
| number of scaffolds | 1,195 | 1,853 | unknown | 14 | 1 |
| max genes per scaffold | 3 | 31 | omega | 10 | 0 |
| *NBS-LRR (Pfam)* | 627 | 1,185 | *Glutenins* |  |  |
| partial genes | 611 | 11 | HMW | 6 | 1 |
| full length genes | 16 | 1,174 | LMW | 16 | 1 |
| number of scaffolds | 613 | 979 | *Prolamins* |  |  |
| max genes per scaffold | 2 | 13 | Avenin | 4 | 0 |
| *CC-NBS-LRR (NLR-parser)* | 225 | 859 | Farinin | 4 | 0 |
|  |  |  | Globulin | 2 | 1 |
|  |  |  | Hordein | 1 | 0 |
|  |  |  | unknown | 1 | 0 |
|  |  |  | Total | 105 | 13 |

#### Gluten genes

Glutens form the major group of grain storage proteins accounting for 10–15% of grain dry weight and confer visco-elastic properties essential for bread-making (Shewry et al., 1995). Gluten genes encode proteins rich in glutamines and prolines that form low complexity sequences composed of PxQ motifs, and occur in tandem repeats in highly complex loci that have posed significant challenges for their assembly and annotation. We characterised the gluten genes in the TGACv1 assembly and showed that most of the known genes were fully assembled. Gluten loci, while still fragmented, exhibit much greater contiguity than in the CSS assembly (IWGSC, 2014) with up to 6 genes per scaffold (Supplementary Figure S12.2). We identified all assembly regions with nucleotide similarity to publicly available gluten sequences, adding an additional 33 gluten genes to the annotation and manually correcting 21 gene models. In total, we identified 105 full length or partial gluten genes and 13 pseudogenes in the TGACv1 assembly (Table 4, Supplementary infomation S12.2).

#### The gibberellin biosynthetic and signalling pathway

Mutations in the gibberellin (GA) biosynthetic and signal transduction pathways have been exploited in wheat, where gain-of-function mutations in the GA signalling protein Rht-1 confer GA insensitivity and a range of dwarfing effects. Most modern wheat cultivars carry semi-dominant *Rht-1* alleles (Phillips, 2016), but these alleles also confer negative pleiotropic effects, including reduced male fertility and grain size. Hence, there is considerable interest in developing alternative dwarfing alleles based on GA-biosynthetic genes such as *GA20ox2*. A prerequisite for this is access to a complete set of genes encoding the biosynthetic pathway. Figure 6 shows that the TGACv1 assembly contains full-length sequences for 67 of the expected 72 GA pathway genes, in contrast to only 23 genes in the CSS assembly (IWGSC, 2014). Two paralogues of *GA20ox3* on chromosome 3D are separated by 460kbp, and *GA1ox-B1* and *GA3ox-B3* are separated by 3.2kbp, suggesting common ancestry of these two enzymes with different catalytic activities (Pearce et al., 2015).

**Figure 6:**
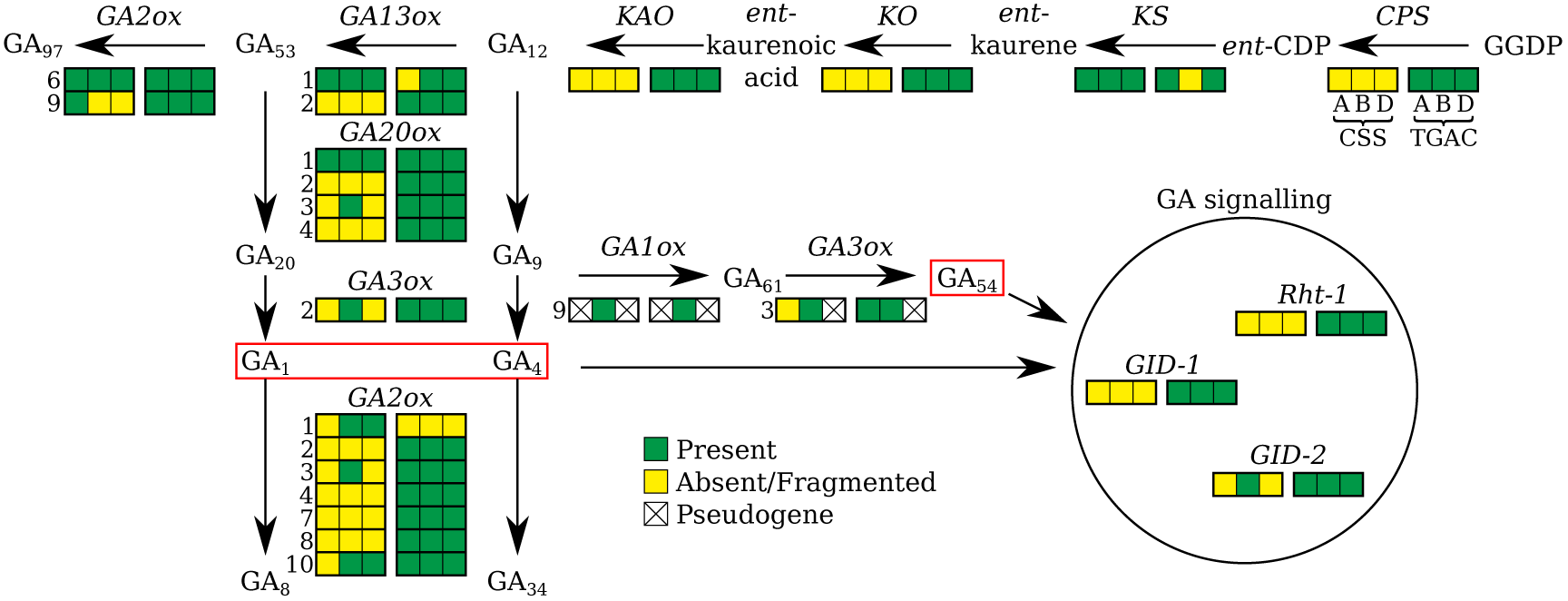
Genes encoding the Gibberellin Biosynthetic and Signalling pathway in bread wheat. The GA biosynthesis, inactivation and signal transduction pathway, illustrating the representation of the gene sequences in CSS and TGACv1 assemblies. If more than one paralog is known for a gene, its number according to the classification by Pearce et al. (2015) is indicated on the left of the box. Bioactive GAs are boxed in red.

## Discussion

Access to a complete and robust wheat genome assembly is essential for the continued improvement of wheat, a staple crop of global significance with 728m tonnes produced in 2014 (http://fenix.fao.org/faostat/beta/en/#home). The capacity to assemble and annotate wheat genomes accurately, rapidly and cost-effectively addresses key social, economic and academic priorities by facilitating trait analyses, by exploiting diverse germplasm resources, and by accelerating plant breeding. However, polyploidy and the extensive repeat structure present in wheat have limited the completeness of previous assembly efforts (Brenchley et al., 2012; IWGSC, 2014; Chapman et al., 2015), reducing their utility.

Here we report the most complete wheat genome assembly to date, representing almost 80% of the 17Gbp genome in large scaffolds. We combined high-quality PCR-free libraries and precisely size-selected LMP libraries (Heavens et al., 2015) with the w2rap assembly software (Clavijo, 2016) to generate contiguous and complete assemblies from relatively low (~33×) Illumina paired-end read coverage and long-mate pair libraries. The contiguity of the TGACv1 assembly allowed us to create a greatly improved gene annotation supported by extensive transcriptome data. Over 78% of the 104,091 high confidence protein coding genes are fully supported by RNA-Seq data. These improvements idenitfied 22,904 genes that were absent from previous wheat gene sets (IWGSC, 2014; Choulet et al., 2014), almost all of which have a homolog in other species (Figure 3B). The robustness of the annotation is further supported by the use of high-quality PacBio data and agreement with proteomic data, with 42% of the HC gene models supported by sequenced peptides. This new wheat gene set provides an improved foundation for wheat research. Finally, incorporation of strand-specific Illumina RNA-Seq libraries into the annotation showed that nearly half of the high confidence genes were alternatively spliced, in line with observations in many other plants (Zhang et al., 2015).

A well-defined gene set in large sequence scaffolds is an essential foundation for trait analyses in wheat. We identified the complement of disease resistance genes, gluten protein genes that confer nutritional and bread-making quality of wheat grains, and the set of gibberellin biosynthetic and signal transduction genes that are important determinants of crop height and yield. An accurate gene set is also essential for understanding expression of gene families in complex allopolyploid genomes. We observed that 20% of homoeologous triads showed differential expression in seedling leaves subject to biotic and abiotic stress conditions. This is consistent with co-expression analyses in developing grains (Pfeifer et al., 2014), where most differentially expressed genes were single homoeologs being up/down-regulated. Taken together, these results identify widespread sub-functionalisation of homoeologous genes due to differential regulation. The new assembly and annotation will enable the identification of multiple sequence differences in promoters, transcription start sites, gene splicing and other features among strict homoeologs, providing a foundation for systematic analyses of the causes of these differences.

Our rapid, accurate and cost-effective assembly approach will enable multiple wheat and Triticeae genomes to be assembled in robust and comparable ways, using relatively inexpensive sequencing technologies and open-source software. These assemblies will reveal a wide spectrum of genetic variation, including large-scale structural changes such as translocations and chromosome additions that are known to play a major role in the adaptation of the wheat crop to different growing environments. By adopting this pan-genomics approach, we will enrich our understanding of complex genome evolution and the plasticity of genome regulation, and empower new approaches to wheat improvement.

## Methods

### DNA library preparation and sequencing

A full description of the DNA preparation and sequencing methods is in Supplemental Information. PCR-free Paired-End (PE) libraries were sequenced using 2×250bp reads on HiSeq2500 platforms for contig generation. Tight, Amplification-free, Large insert Libraries (TALL) libraries and Nextera LMP libraries (Heavens et al., 2015) were used for scaffolding. Insert size distributions (Supplemental Figures S4.1, S4.3 and S4.2) were checked by mapping to the CS42 chromosome 3B pseudo-molecule (Choulet et al., 2014) using the DRAGEN co-processor (http://www.edicogenome.com/dragen/).

### Assembly

Assembly was performed using the Wheat/Whole Genome Robust Assembly Pipeline, w2rap (Clavijo, 2016). It combines the w2rap-contigger, based on DISCOVAR *de novo* (Weisenfeld et al., 2014), an LMP preparation approach based on FLASH (Magoc and Salzberg, 2011) and Nextclip (Leggett et al., 2014), and scaffolding with SOAPdenovo2 (Luo et al., 2012). The w2rap-contigger takes advantage of DISCOVAR (Weisenfeld et al., 2014; Love et al., 2016) algorithms to preserve sequence variation during assembly, but has been further developed to enable processing of much larger data volumes and complex genomic repeats. The paired end read dataset was assembled into contigs on a SGI UV200 machine with 64 cores and 7TB of shared RAM, using the default settings of the w2rap-contigger from https://github.com/bioinfologics/ w2rap-contigger/releases/tag/CS42_TGACv1. Contigs were scaffolded using the PE, LMP and TALL reads and the SOAPdenovo2 (Luo et al., 2012) prepare->map->scaffold pipeline, run at k=71 for both the prepare and map steps on the same machine using 128 cores. Contigs and scaffolds were QC’ed using KAT spectra-cn plots (Mapleson et al., 2016a) to assess motif representation.

### Gene annotation

A high quality gene set for wheat was generated using a custom pipeline integrating wheat-specific transcriptomic data, protein similarity, and evidence-guided gene predictions generated with AUGUSTUS (Stanke and Morgenstern, 2005). Full methods are in Supplementary Information 8. RNA-Seq reads (ERP004714, ERP004505, and 250bp paired-end strand-specific reads from six different tissues) were assembled using four alternative assembly methods (Trapnell et al., 2010; Pertea et al., 2015; Song et al., 2016; Haas et al., 2013) and integrated with PacBio transcripts into a coherent and non-redundant set of models using Mikado (Venturini et al., 2016). PacBio reads were then classified according to protein similarity and a subset of high quality (e.g. full length, canonical splicing, non-redundant) transcripts used to train an AUGUSTUS wheat-specific gene prediction model. AUGUSTUS was then used to generate a first draft of the genome annotation, using as input Mikado-filtered transcript models, reliable junctions identified with Portcullis (Mapleson et al., 2016b), and peptide alignments of proteins from five close wheat relatives (*B. distachyon*, maize, rice, *S. bicolor* and *S. italica*). This draft annotation was refined by correcting probable gene fusions, missing loci and alternative splice variants. The annotation was functionally annotated and all loci were assigned a confidence rank based on their similarity to known proteins and their agreement with transcriptome data.

## Data access

All data generated in this study has been submitted to the EMBL-EBI European Nucleotide Archive. PE and LMP reads used for genome assembly and scaffolding are available in study accession PRJEB15378, and assembled scaffolds are available in study accession PRJEB11773. The Illumina and PacBio reads used for genome annotation are available in study accession PRJEB15048. The assembly and annotation is available in Ensembl Plants (release 32) at http://plants.ensembl.org/Triticum_aestivum and from the Earlham Institute server at http://opendata.earlham.ac.uk/Triticum_. BLAST services for these datasets are available at https://wheatis.tgac.ac.uk/grassroots-portal/blast.

## Acknowledgements

We thank Burkhard Steuernagel for assistance with NLR-parser. This work was funded by a Biotechnology and Biological Sciences Research Council (BBSRC) strategic LOLA Award to MWB and CU (BB/J003557/1), MDC (BB/J003743/1), PK (BB/J00328X/1) and AP (BB/J003913/1), a BBSRC Anniversary Future Leader Fellowship (BB/M014045/1) to PB, BBSRC Institute Strategic Programme Grants GRO (BB/J004588/1) to MWB and CU, “2020 Wheat” (BBS/E/C/00005202) to AP, and Bioinformatics BB/J004669/1 to FDP. The German Ministry of Education and Research (BMBF) grant 031A536 “de.NBI” supported MS, and the Australian Research Council (LP120200102, CE140100008) and Agilent Technologies Australia supported AHM. Next-generation sequencing and library construction was delivered via the BBSRC National Capability in Genomics (BB/J010375/1) at the Earlham Institute (EI, formerly The Genome Analysis Centre, Norwich), by members of the Platforms and Pipelines Group. Open data access, and BLAST databases and service, are provided by the EI Data Infrastructure group.

